# Deep Learning Enables Individual Xenograft Cell Classification in Histological Images by Analysis of Contextual Features

**DOI:** 10.1101/2020.11.03.361741

**Authors:** Quentin Juppet, Fabio De Martino, Martin Weigert, Olivier Burri, Michaël Unser, Cathrin Brisken, Daniel Sage

## Abstract

Patient-Derived Xenografts (PDXs) are the preclinical models which best recapitulate inter- and intra-patient complexity of human breast malignancies, and are also emerging as useful tools to study the normal breast epithelium. However, data analysis generated with such models is often confounded by the presence of host cells and can give rise to data misinterpretation. For instance, it is important to discriminate between xenografted and host cells in histological sections prior to performing immunostainings. We developed Single Cell Classifier (SCC), a data-driven deep learning-based computational tool that provides an innovative approach for automated cell species discrimination based on a multi-step process entailing nuclei segmentation and single cell classification. We show that human and murine cells contextual features, more than cell-intrinsic ones, can be exploited to discriminate between cell species in both normal and malignant tissues, yielding up to 96% classification accuracy. SCC will facilitate the interpretation of H&E stained histological sections of xenografted human-in-mouse tissues and it is open to new in-house built models for further applications. SCC is released as an open-source plugin in ImageJ/Fiji available at the following link: https://github.com/Biomedical-Imaging-Group/SingleCellClassifier.

**Author summary:** Breast cancer is the most commonly diagnosed tumor in women worldwide and its incidence in the population is increasing over time. Because our understanding of such disease has been hampered by the lack of adequate human preclinical model, efforts have been made in order to develop better approaches to model the human complexity. Recent advances in this regard were achieved with Patient-Derived Xenografts (PDXs), which entail the implantation of human-derived specimens to recipient immunosuppressed mice and are, thus far, the preclinical system best recapitulating the heterogeneity of both normal and malignant human tissues. However, histological analyses of the resulting tissues are usually confounded by the presence of cells of different species. To circumvent this hurdle and to facilitate the discrimination of human and murine cells in xenografted samples, we developed Single Cell Classifier (SCC), a deep learning-based open-source software, available as a plugin in ImageJ/Fiji, performing automated species classification of individual cells in H&E stained sections. We show that SCC can reach up to 96% classification accuracy to classify cells of different species mainly leveraging on their contextual features in both normal and tumor PDXs. SCC will improve and automate histological analyses of human-in-mouse xenografts and is open to new in-house built models for further classification tasks and applications in image analysis.

## Introduction

Most of our understanding of mammary gland development and breast carcinogenesis stems from experiments with animal models. Mice are by far the most widely used experimental system due to their size, ease of use, and most importantly, because of the many powerful genetic tools which are available for this species. However, approximately 90% of potential oncology drugs fail in clinical trials [1, 2], partly because of the lack of adequate preclinical models, raising concerns on how representative of the human physiology and disease data derived from mice is.

Patient-Derived Xenografts (PDXs), namely preclinical models developed by transplanting human-derived cells into immunosuppressed or humanized mice, currently best recapitulate the complexity of human tissues and are increasingly employed for translational research [3–5]. Classically, mammary xenografts are obtained by orthotopical transplantation of pieces of primary breast tissues to the mammary fat pad of recipient immunosuppressed mice [6]. However, under these settings the PDXs growth and their HR expression are dependent on estradiol supplementations which, resulting in serum E2 equivalent to mid-menstrual cycle levels [6, 7], alter the physiological relevance of this preclinical system. Recent advances in this field were achieved with the Mouse INtraDuctal (MIND) model, which entails the injection of primary human-derived breast cells directly into the mouse mammary ductal tree *via* cleaved teat [8, 9]. In the intraductal microenvironment primary HBECs and breast cancer cells grow independently of any hormone supplementations while retaining their HR expression and hormone responsiveness, making the MIND model an appealing preclinical tool [8, 10–14]. Hence, the MIND model provides the unprecedented opportunity to study the role that individual HRs play in the luminal compartment of the human breast epithelium by, for instance, histological techniques [9]. However, molecular analyses in xenograft models are hindered by the presence of both human and murine cells, which could possibly lead to data misinterpretation due to contamination of different cell species. Therefore, prior to performing specific immunostaining to assess levels of proteins of interest in the xenografted cells, paraformaldehyde-fixed paraffin-embedded sections are usually stained by Haematoxylin and Eosin (H&E) in order to carry out a rough inference of the abundance of human cells within the tissue of interest based on morphological features. Although human cells are usually bigger in size and more elongated than their murine counterparts, manual evaluation is usually error-prone, time-consuming and subject to inter-personal variability.

Machine learning techniques have been effectively applied in a number of different fields and emerged as valuable resources to decipher the content of biological images [15, 16]. While some methods have already been developed to analyze H&E stained human histological sections, their aim was mainly tissue segmentation [17–19] and more sophisticated supervised learning-based tools have been created to perform nuclei segmentation [19]. Here, we hypothesized that deep learning could be employed in order to automate human-mouse cell discrimination in intraductal xenografts, a challenging task due to high biological and technical heterogeneity. We developed Single Cell Classifier (SCC), a data-driven machine learning-based approach capable of classifying either normal or malignant individual xenografted cells from murine cells in the same histological section according to specific features rather than images. Upon evaluation of a total of 484 cell-intrinsic or contextual features, SCC was proven to reach up to 96% of classification accuracy, with contextual information playing a major role on the classification performance. SCC is supplied as a publicly available plugin in ImageJ/Fiji [20] and can be downloaded at the following link: https://github.com/Biomedical-Imaging-Group/SingleCellClassifier.

## Materials and methods

### Intraductal Injections

Single cells readily isolated from reduction mammoplasty specimens or invasive breast cancers were lentivirally infected with firefly luciferase (luc2) and GFP (Lenti-ONE CMV-GFP(2A)Luc2 WPRE VSV, ref = V621004001 or pR980 Luc2GFP, GEG Tech), allowing to track the growth of HBECs *in vivo*, and promptly xenografted into 7-to-12 week old NOD.Cg-Prkdcscid Il2rgtm1Wjl/SzJ (NSG) mice. Animals were anesthetized by intraperitoneal injection with 10 mg*/*kg xylazine and 90 mg*/*kg ketamine (Graeub) and intraductally injected with 10 µL of PBS containing 500.000 cells per gland, as previously described [9]. At sacrifice, xenografted mammary glands were harvested for histological analyses.

### Histology

Tissues were fixed in 4% paraformaldehyde and paraffin-embedded for histological analyses. Paraffin blocks were cut and 4 µm-thick sections were mounted onto 75×25 mm Superfrost Plus microscope slides (Thermo Scientific, USA, ref = J1800AMNZ). H&E staining was performed according to standard protocol. For immunostaining, sections were deparaffinized in xylene and re-hydrated. Antigen retrieval was carried out in 10 mmol sodium citrate (pH 6.0) at 95 ^*°*^C for 25 min. Blocking was performed with 1% BSA for 60 minutes. Sections were incubated overnight with primary antibodies, followed by 1-hour incubation with secondary antibodies. For fluorescence microscopy, nuclei were counterstained with DAPI (Sigma) and then mounted with Fluoromount-GTM (cat# 4958-02, Invitrogen). E-cadherin antibody: clone G10; sc-8426; dilution 1:100; Santa Cruz. Cytokeratin 7 antibody: clone SP52; ab183344; dilution 1:500; Abcam. Secondary anti-mouse antibody: Alexa 488-conjugated; clone A-11029; dilution 1:700; Thermo Fisher Scientific.

### Image Acquisition

Slides were scanned with Olympus VS120-L100 slide scanner using a 20x/0.75 objective connected to a Pike F505 C Color camera. Because of the size of the resulting data (Fig 1), images were loaded into QuPath [21] using the BioFormats extension (https://github.com/qupath/qupath-bioformats-extension). The slide images are publicly available on Zenodo (https://zenodo.org/record/3960270).

**Fig 1.**
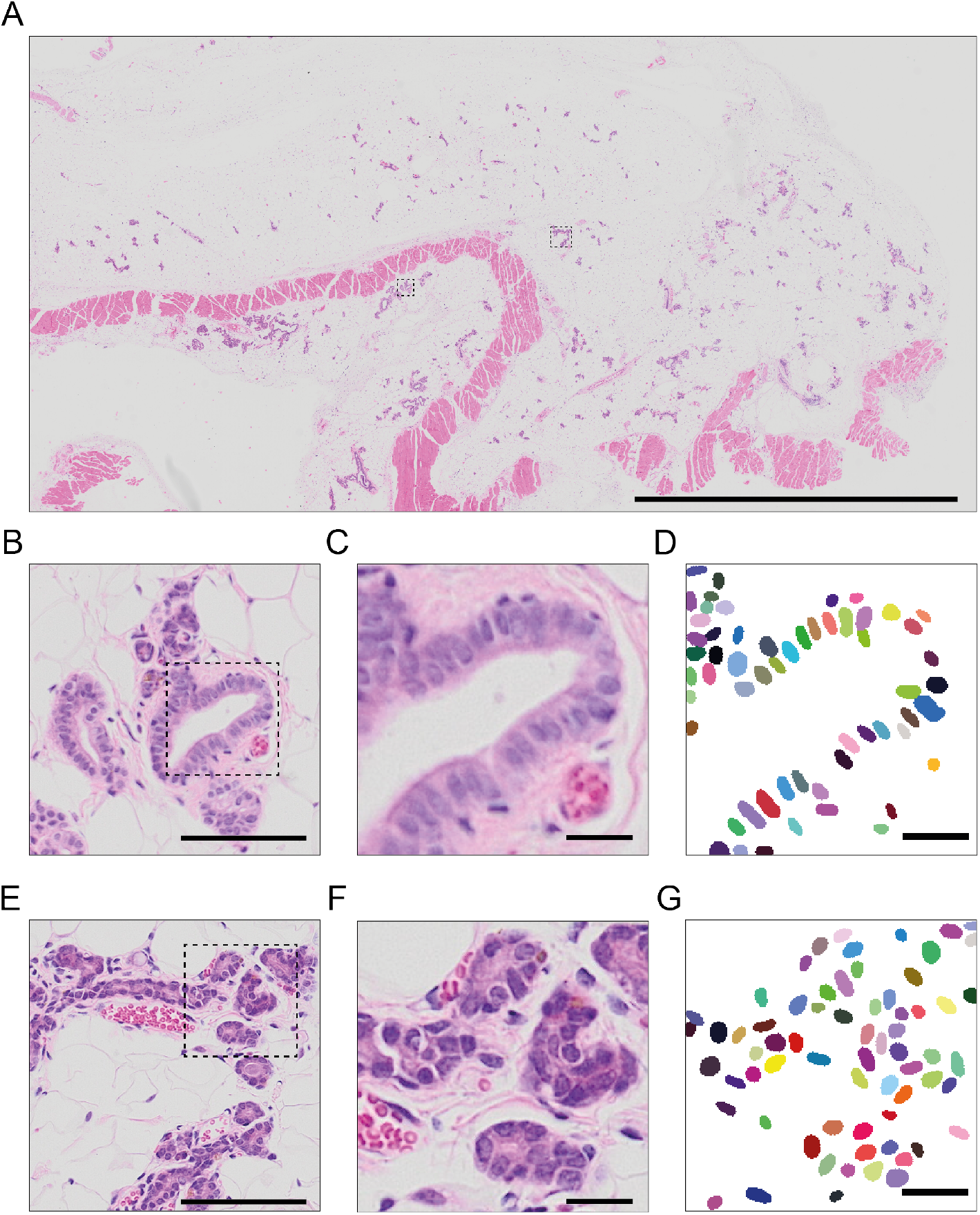
Characterization of histological images of H&E-stained mouse mammary gland intraductally xenografted with human breast cells. A: Complete slide image, image size of approximately 1 billion pixels. Boxed areas mark human and mouse cell clusters. Scale bar, 5000 µm. B: Magnified area of a human cluster. Scale bar, 100 µm. C: Magnified area of human cells. Scale bar, 20 µm. D: Manual annotations of the human nuclei associated with the cells randomly colored for distinction purpose. Scale bar, 20 µm. E: Magnified area of a murine cluster. Scale bar, 100 µm. F: Magnified area of murine cells. Scale bar, 20 µm. G: Manual annotations of the murine nuclei associated with the cells randomly colored for distinction purpose. Scale bar, 20 µm.

### Data Extraction

Due to the pyramidal nature of the whole slide scanner images, a version of each image was extracted to define areas likely to contain ducts. Because ducts can be defined as relatively large, densely packed cell regions having a dense Eosin signal, we can use this information to extract them using Fiji’s [20] Color Deconvolution with the built-in “H&E DAB” vectors. First, a 4-fold downsampled version of each whole slide image was sent from QuPath [21] to ImageJ [20]. The extracted signal was then filtered with a Gaussian kernel of *σ* = 2 px before thresholding with ImageJ’s [20] Default method. Finally, connected components analysis (AnalyzeParticles) was used to obtain ROIs. The bounding boxes of these ROIs were extracted and enlarged slightly to ensure no ducts were touching the edge of the image. Resulting bounding boxes were reimported into QuPath [21] as annotations and used to perform the export of the full resolution ducts as .tif images.

### Nuclei Detection

As defining precise boundaries between neighboring cells can be challenging, we segment the nuclei as a first step using StarDist [22, 23], a state-of-the-art method outperforming classical approaches to detect star-convex objects for 2D images using a neural network. Such networks can be trained from various image types and the detection is represented as a labeled image where each nucleus is associated with a label.

A StarDist model was trained with a set of 24 images of size 320×320 pixels extracted from H&E-stained sections of normal human breast xenografted mammary glands (Sup. Fig 1). The training data is publicly available on the GitHub of SCC. The nuclei of these images were manually annotated for a total of approximately 2500 nuclei. The training was performed in Python using TensorFlow on Google Colab with GPU on 100 epochs for 40 minutes. The StarDist model was configured to detect 48 rays objects and a dropout of 0.5 was added to the network to avoid overfitting.

### Cells Delineation

The estimation of the cells (Fig 2F) was computed according to two criteria: (i) cells should not overlap with each other and (ii) their thickness, defined as distance to the nuclei, should not exceed Δ = 2 µm. This maximal value is an estimation of the expected thickness for the type of cells we aim to classify, and can be edited by the users. The mask of the first criterion was computed using a Voronoi diagram on the nuclei labels and of the second criterion by thresholding, at Δ, a distance map of the nuclei labels. Hence, each detected cell will be associated with its nucleus. These criteria can be represented as masks that can be integrated to delineate each cell.

**Fig 2.**
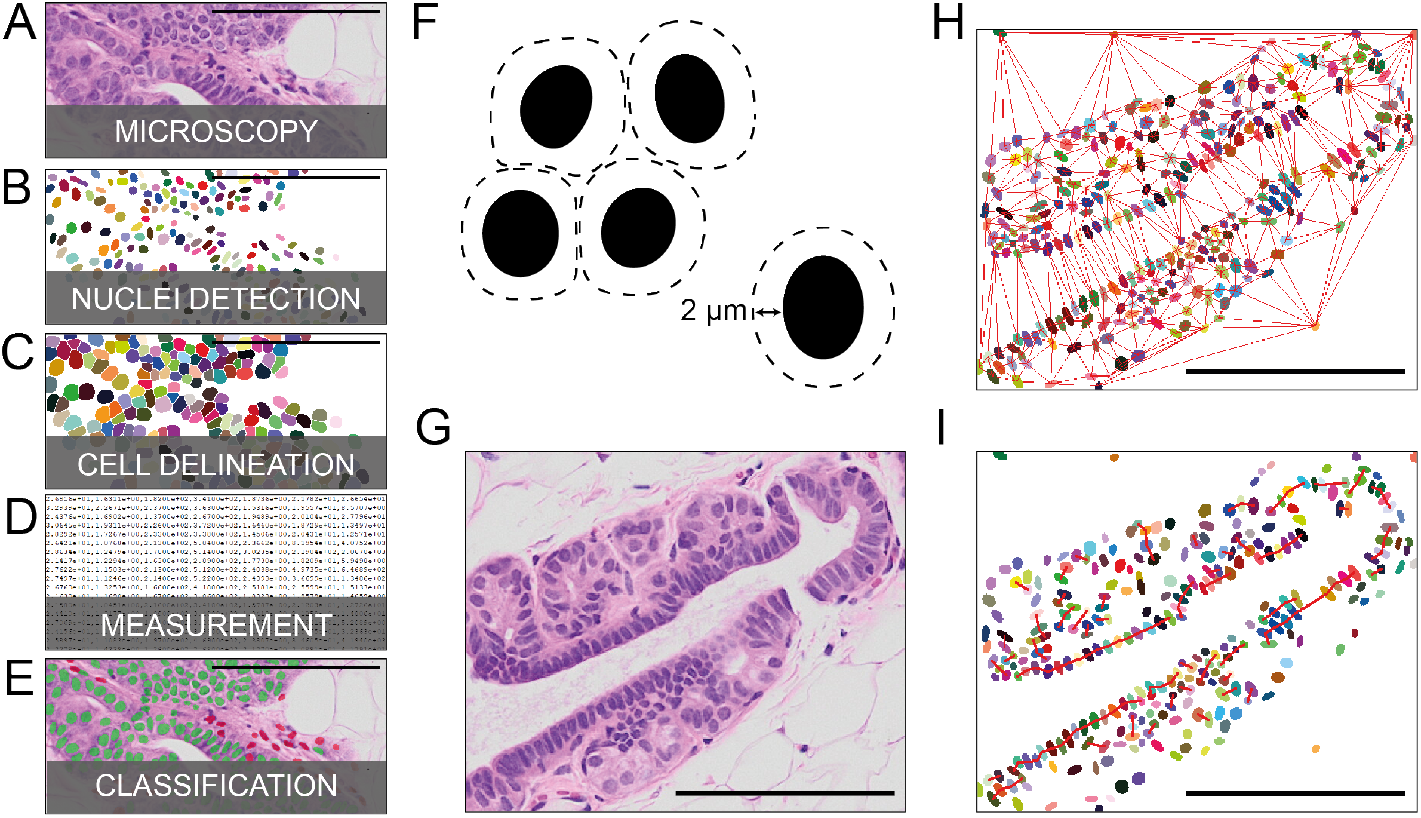
SCC pipeline. A: Source example for the pipeline, an H&E-stained “humanized” mouse mammary duct. B: Label image of the nuclei detected by StarDist randomly colored for distinction purposes. C: Label image of the cells estimated from the nuclei randomly colored for distinction purpose. D: Features are extracted from the nuclei and the cells. E: Input image overlaid with the cell class computed from their associated features with a neural network. Green color defines human cells and red murine cells. F: Diagram of the cell delineation with respect to two criteria: cells should not overlap with each other and the cell thickness (distance to the nuclei) should not exceed 2 µm. G: Source example for the contextual features, an H&E-stained “humanized” mouse mammary duct. H: The Delaunay graph in red over the detected nuclei associated with the source, randomly colored for distinction purpose. I: The cell chain graph over the detected nuclei associated with the source, randomly colored for distinction purpose, red segments correspond to cell chain. Scale bars, 100 µm.

### Measurement

To classify cells, it is crucial to properly describe them. To do so, 484 features were extracted from the detected nuclei and cells. These features can be related to the object itself (i.e. cell-intrinsic features), or related to their neighbors and their organization (i.e. contextual features). Out of these, 47 concern cell-intrinsic features, 376 contextual features are related to the cell-intrinsic features of the neighboring cells and the remaining 61 contextual features describe the organization of the neighbors.

### Shape and Size Features

Cell-intrinsic features related to the shape and the size of both nuclei and cells were measured by fitting an ellipse on the targeted object to extract its elongation, minor axis, and major axis as features. Also, the area, and in particular the area ratio between the nuclei and the cell, were exploited for the analysis.

### Textural Features

Cell-intrinsic textural features were extracted from the pixels of the source images at specific areas for each cell. The cells were divided into two areas: (i) their nuclei and (ii) their estimated cytoplasms, defined as the cell area without the nucleus. Such features can be divided between features related to the color and features related to the texture itself. For the color-related features, the mean and the variation of the values of each channel were computed. For the features related to the texture, the Haralick texture features [24, 25] were calculated on a gray level version of the source image. The Haralick texture features provide 13 statistical parameters related to the pixels such as their entropy or their contrast. The gray level image was computed from a principal component analysis (PCA) on the channels of the image, with the first component corresponding to the factors to apply on the channels, thereby maximizing their variance, so that the maximum amount of information can be measured for the texture.

### Contextual Features

In addition to cell-intrinsic features, we acquired contextual features. Indeed, cells have many ways to spatially organize themselves and, moreover, neighboring cells can share peculiar features.

To efficiently determine the closest neighbors of a cell, a Delaunay graph [26] (Fig 2H) was computed, on which a Dijkstra algorithm was employed. The cells are represented as points on this graph, with each point corresponding to the centroids of the cells.

For each cell, many kinds of neighboring structures were inspected, providing specific features:

i. The neighbors directly connected to the cell on the Delaunay graph give information about the position of a given cell in the cluster given the mean and the variance of their distance. A cell described by high distance variance from its direct neighbors will have a high probability to be located at the border of a cluster.
ii. The lateral neighbors are the two cells located on the left and the right of a given cell given the orientation of the nucleus ellipse. Features like the alignment, the distance, and the difference in orientation were measured. When two cells are mutually lateral neighbors, they can be iteratively connected to create a chain of cells (Fig 2I). Features related to these chains, which are relatively abundant in human cell clusters, are their tortuosity and size.
iii. To cover different types of cell aggregates, 4 sizes of vicinity were analyzed represented by *K* equal to 5, 10, 20, and 40 cells. These *K* neighbors are the set of the *K* closest neighbors to the current cell. Such distribution allows the extraction, at the same time, of both local and global features depending on the *K*. It is possible to extract distance features (mean and variance) between the *K* neighbors and the current cells and, in particular, the distance of the current cell to the centroid of the set of *K* neighbors. The distance between the current cell and the centroid of the set provides information about the homogeneity in the set since, in a homogeneous cluster, it is expected that the current cell is very close to the position of the centroid as the neighbors should be spread around the current cell homogeneously.
iv. The *K* connected neighbors are the set of the *K* closest neighbors to the cells that are physically connected to each other. Like the *K* neighbors, the *K* connected neighbors provide at the same time both local and global information of the neighborhood of the cell taken into account, by considering the same *K* values. Moreover, a peculiar evidence is that cells that are physically connected share common features. This property motivated the extraction of the cell-intrinsic features of the neighbors through means and variances. It is also relevant to observe the shape of the cluster formed by the neighbors, which can be performed by computing an ellipse in a similar fashion than what was previously performed for the nuclei and cells shape features.

### Classification

To classify the cells based on their features, a neural network model was trained by supervised learning in Python using TensorFlow. A set of 174 images of various sizes were extracted from H&E-stained xenografted mouse mammary glands (Sup. Fig 2). Among these images, 96 mostly contain normal human cells, whereas 78 contain only mouse cells. To ensure that an image contains normal human cells, a human-specific E-cadherin (E-CAD) antibody was used in order to uniquely probe for xenografted non-malignant cells. This set of images represented about 60.000 cells after detection, of which are human. Thereby, 26.000 mouse cells were considered to balance the number of cells between the two classes of interest such that their influence on the training is equivalent.

To classify tumor-derived PDXs, 17 images that contain mostly tumor human cells were added to the previous set. In order to circumvent interpretation problems due to dynamic E-cadherin protein expression levels in tumor cells [27], tumor cells were uniquely detected exploiting a human-specific cytokeratin 7 (CK7) antibody. The final set represented about 80.000 cells with about 40.000 mouse cells, 26.000 normal human cells, and 14.000 tumor human cells.

To define the species of a cell, classes masks were manually annotated for each of the images based on their fluorescent control.

The design of this network contains 4 fully connected layers with a decreasing number of neurons: 128, 64, 32, and 16 neurons. Each layer is associated with a dropout of 0.5 and batch normalization to avoid overfitting. Their activation function is a rectified linear unit (ReLU). At the end of the network, the output layer is a fully-connected layer with 2 neurons corresponding to the two classes, namely human and mouse, associated with a sigmoid activation function. The training minimizes a categorical cross-entropy loss and was optimized using the Adam optimization algorithm, which takes as input the 484 features and returns the probability of a cell to belong to both human or mouse class as output.

### ImageJ/Fiji Plugin

An ImageJ/Fiji [20] plugin named “Single Cell Classifier” has been implemented to allow users to perform cell classification using our method. It also provides the possibility to adopt other models than the built-in ones in order to allow for the classification of other classes of objects, other types of tissues or other types of images, such as fluorescence-based ones. The parameters of the methods such as the K values for the neighbors, the factors to convert in gray value, or the cell thickness Δ are also editable by the user.

This plugins depend on three other plugins: StarDist [22, 23] for the nuclei detection, MorphoLibJ [28] for its morphological and analysis tools used in cells delineation and features extraction, and finally CSBDeep [29] that execute our classification neural network.

## Results

### Nuclei Detection

The nuclei detection performed by our StarDist model reaches an accuracy of 74.74% (Table 1), remaining accurate to distinguish nuclei even under challenging conditions (Fig 3).

**Table 1.** Segmentation result statistics.

|  |  |
| --- | --- |
| Accuracy | 74.74% |
| Precision | 86.52% |
| Sensitivity | 84.59% |
| F<sub>1</sub> score | 85.54% |

**Fig 3.**
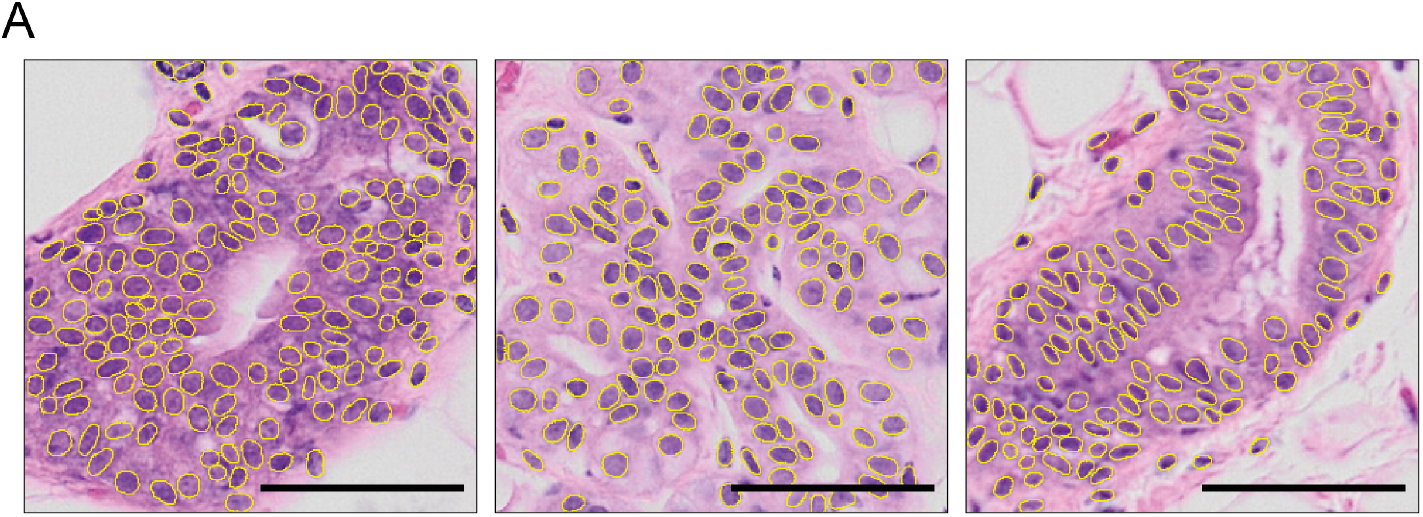
Nuclei segmentation performance. Examples of nuclei segmentation, with detection outlined in yellow. Scale bars, 50 µm.

### Features Analysis

Next, we performed a comparison between human and mouse features to characterize the impact of each feature on the neural network to assess the morphology of the engrafted human cells in the intraductal environment. The analysis of the features distribution and their correlation with the classes revealed that shape and size of individual cells poorly discriminate the two classes, with a correlation index lower than 0.2, suggesting that they do not help the discrimination task (Fig 4A, 4B). However, the contextual features for shape and size have a better correlation ranging between 0.3 and 0.4, highlighting the importance of the context for this purpose (Fig 4A, 4C).

**Fig 4.**
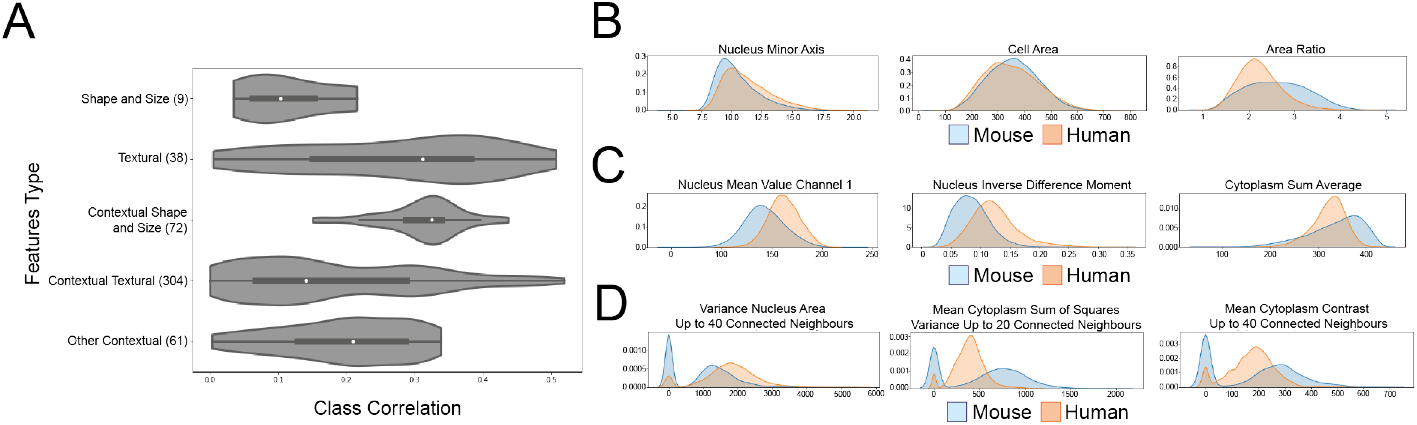
Features analysis. A: Violin plot of the correlation between the features, grouped by categories, and the classes. The features number in each category is indicated in parenthesis. The other contextual features refer to the contextual features related to the organization of the neighbors and not related to cell-intrinsic features. B: Kernel density estimation plot of some shape and size features. C: Kernel density estimation plot of some textural features. D: Kernel density estimation plot of some contextual features. The value 0 is assigned to cells that do not have enough neighbours.

Interestingly, textural features seem to offer better help in the cell species discrimination than shape and size, as highlighted by the resulting small degree of overlap between the two cell species and their global better correlation with the classes, concentrated between 0.15 and 0.35 but reaching a maximum of 0.5. The context seems to decrease the correlation for most of the features, concentrated between 0.05 and 0.3 (Fig 4A, 4D). However, some exceptions in the contextual texture features still reach a high correlation of 0.5, making them appealing for such analysis.

### Classification of Normal Breast PDXs

SCC performs normal human breast cell discrimination reaching an accuracy of 96.51% (Fig 5) as assessed by quantifying the number of accurate calls upon manual annotation of humanized mouse mammary ducts. Both sensitivity and precision are higher than 96% for the classification of both classes taken into consideration (Table 2). The Area Under the Curve of the Receiver Operating Characteristics (AUC ROC) was used as a standard to estimate our predictive power and revealed that SCC efficiently predicts both human and mouse cells with probabilities associated with each of the analyzed cells close to the extrema 0 and 1, suggesting high confidence of our model. Interestingly, the feature importance previously discussed in Section impinges on the classification task (Table 3). In line with our predictions, we observed that the contextual features without any texture information allow for a very good classification with 89.02% of accuracy, compared to the shape and size features alone achieving an accuracy of 66.84%, highlighting the importance of contextual features for such classification task.

**Fig 5.**
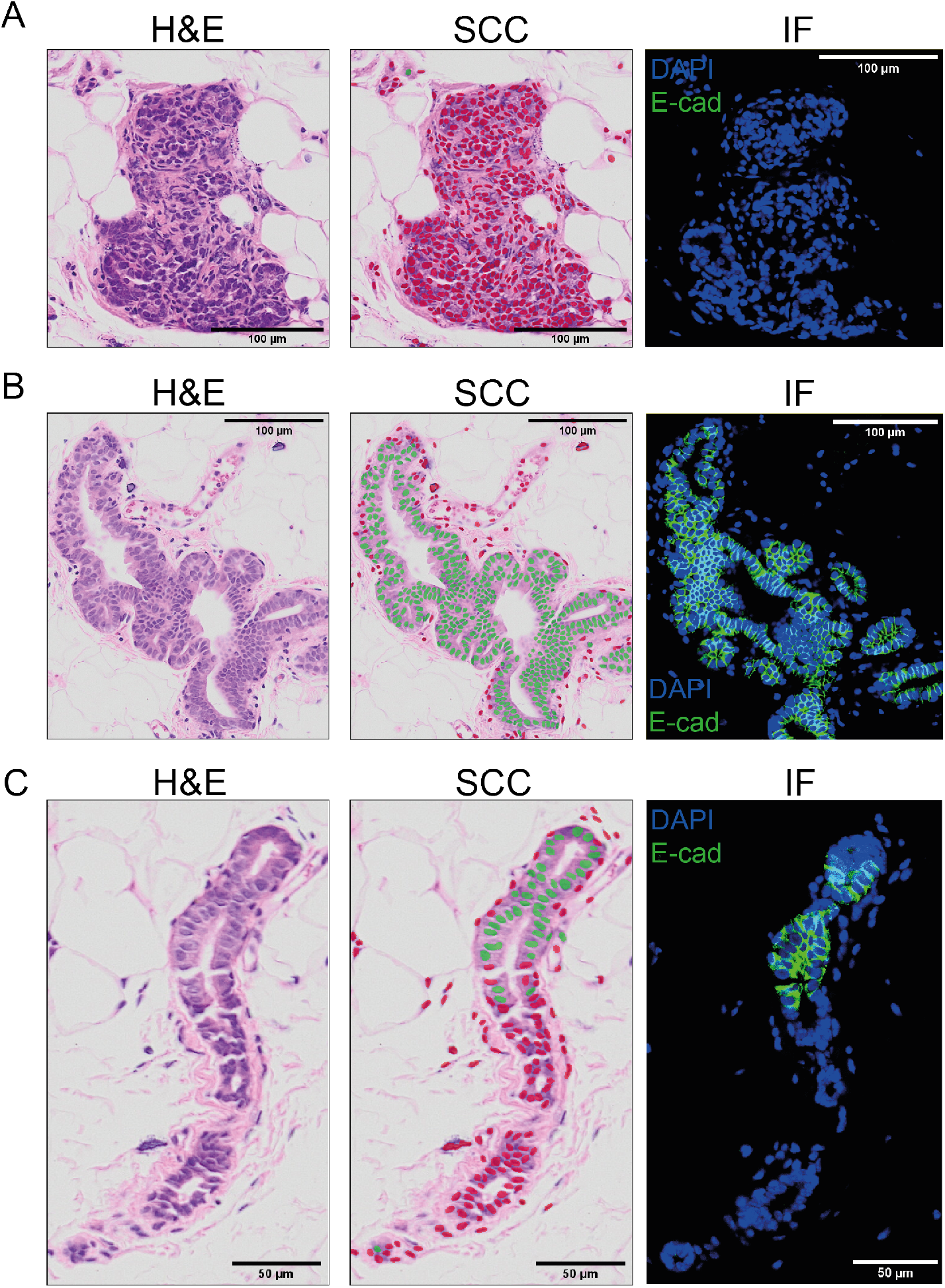
Cell species classification of normal human breast-derived PDXs. A: Example of a cluster of human cells. Green and red dots correspond to human and mouse class, respectively. For the fluorescent controls, the green signal corresponds to a human-specific E-cadherin, blue signal corresponds to DAPI. The source images are on the left, the classification result in the middle and the fluorescent control on the right. B: Example of a cluster of murine cells. Green and red dots correspond to human and mouse class, respectively. For the fluorescent controls, the green signal corresponds to a human-specific E-cadherin, blue signal corresponds to DAPI. The source images are on the left, the classification result in the middle and the fluorescent control on the right. C: Example of a cluster containing both murine and human cells. Green and red dots correspond to human and mouse class, respectively. For the fluorescent controls, the green signal corresponds to a human-specific E-cadherin, blue signal corresponds to DAPI. The source images are on the left, the classification result in the middle and the fluorescent control on the right.

**Table 2.** Classification result statistics.

|  |  |
| --- | --- |
| Accuracy | 96.51% |
| Mouse precision | 96.17% |
| Mouse sensitivity | 96.91% |
| Mouse $F_1$ score | 96.54% |
| Mouse AUC - ROC | 99.48% |
| Human precision | 96.86% |
| Human sensitivity | 96.10% |
| Human $F_1$ score | 96.48% |
| Human AUC - ROC | 99.48% |

**Table 3.** Features impact on classifier accuracy.

|  |  | Contextual |  |
| --- | --- | --- | --- |
|  |  | Without | With |
| Textural | Without | 66.84% ± 0.71% | 89.02% ± 0.54% |
| | With | 85.68% ± 0.33% | 96.41% ± 0.21% |

### Classification of Breast Cancer-Derived PDXs

As patient-derived xenografts are mainly used in the context of cancer research, we went on to assess whether SCC was also able to classify cell species on tumor-derived PDXs. Whereas the segmentation model remained unchanged compared to the one developed for normal human breast cells, we prepared a new classification model in order to discriminate human cancer cells from mouse cells by integrating new tumor intraductal xenografts-derived images to the previous model. Estimation of accuracy in different histological subtypes of breast cancer based on 20 images representing a range of 5000 to 10000 cells per type revealed that SCC successfully classified more than 90% of the cells across different breast cancer subtypes (Table 4). Overall, this new model reaches an accuracy of 96.21% across different breast tumor types (Fig 6). The analysis of how the features impinged on the classification was performed and showed that the textural features have a higher impact compared to the normal human cells, reaching an accuracy of 95.50% (Table 5).

**Table 4.** Tumor type classification accuracies.

| Tumor type | Accuracy |
| --- | --- |
| Invasive Lobular Carcinoma | 94.40% |
| No Special Subtype | 92.62% |

**Fig 6.**
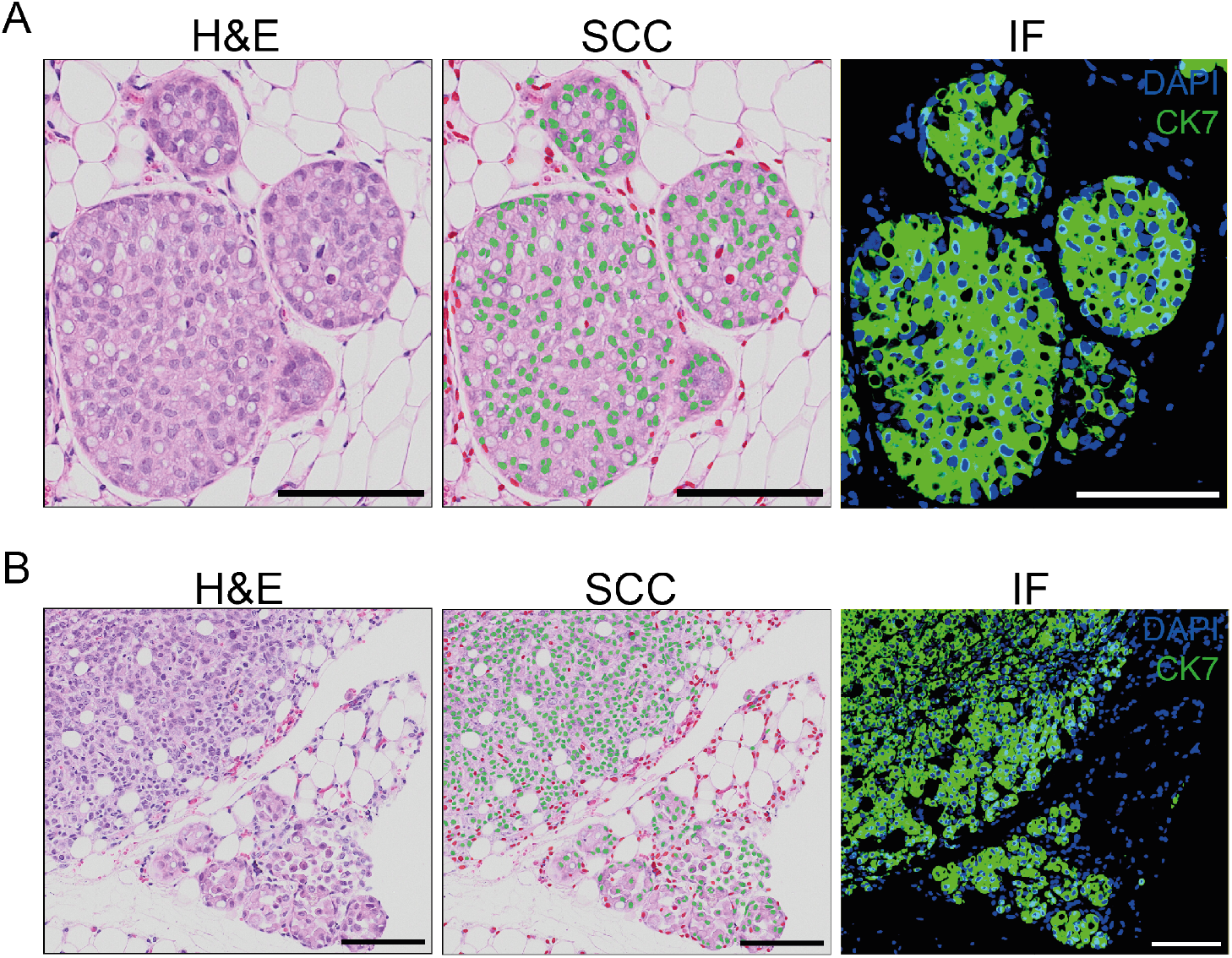
Cell species classification results on breast cancer-derived PDXs. A: Cluster of tumor human cells: source image (left), with its classification result (middle) and its fluorescent control (right). Green and red dots correspond to human and mouse, respectively. For the fluorescent controls, the green signal corresponds to a human-specific CK7, blue signal corresponds to DAPI. B: Cluster of tumor human cells: source image (left), with its classification result (middle) and its fluorescent control (right). Green and red dots correspond to human and mouse, respectively. For the fluorescent controls, the green signal corresponds to a human-specific CK7, blue signal corresponds to DAPI. Scale bars, 100 µm.

**Table 5.** Features impact on classifier accuracy for human tumor cells.

|  |  | Contextual |  |
| --- | --- | --- | --- |
|  |  | Without | With |
| Textural | Without | 65.47% ± 0.42% | 88.65% ± 0.16% |
| | With | 95.50% ± 0.20% | 97.26% ± 0.18% |

### Comparison with an Image-Based Single-Stage Method

Finally, we investigated whether both the nucleus segmentation task and cell classification task could be performed jointly by a single-stage model. To that end, we extended the model architecture of StarDist [22] and added a dedicated classification head that predicts the probability of a nucleus belonging either to a human or a mouse cell. We then annotated nucleus outlines and cell types (human/mouse) for 16 images of size 330×320 containing in total 1450 human and 300 mouse nuclei. After training, the extended model achieved a classification accuracy of 85.9 % across all matched nucleus instances, well below the accuracy achieved by SCC. This suggests that decoupling the segmentation and classification tasks is beneficial in our case possibly due to the availability of only few annotated training data, and that SCC outperforms standard image-based single-stage methods.

## Discussion

PDXs are innovative preclinical models having applications for translational research. However, their usage is hampered by difficulty in data interpretation arising from the presence of host cells and no methods are currently available for species discrimination of individual cells in PDX models. To tackle this lack we developed SCC, a publicly available deep learning-based tool aiming at classifying human and mouse cells in PDX-derived histological sections. For the first time, SCC elaborates a comprehensive set of information to efficiently classify the species of individual cells based on both cell-intrinsic and contextual features proper of the neighboring tissue context. Surprisingly, features analysis revealed that contextual features were more important than cell-intrinsic ones for an accurate cell classification, suggesting that xenografted human cells retain their typical morphologies while creating clusters characterized by distinctive conformations in the host. Although SCC was initially designed for detection of normal breast epithelial cells, we show that the completeness of the set of features taken into consideration in our analysis yielded, on average, up to 96% of discrimination accuracy for both normal and malignant xenografted cells derived from different breast cancer subtypes. The automated classification of individual human and mouse cells performed by SCC can facilitate the work of researchers dealing with PDXs-derived histological sections, making the subsequent image analyses faster and more reproducible, and will enable quantification of xenografted cells to assess grafting efficiency or cell growth. Finally, SCC is a dynamic software as users are allowed to input their in-house models in order to perform classifications between any cells of interest, making it appealing for further applications in the field of image analysis.

## Declarations

## Acknowledgments

The authors wish to acknowledge the support of the Phenogenomics Center, Histology and the Bio Imaging & Optics Core Facility at EPFL for technical assistance, G.Sflomos for reading of the manuscript, A. Ayyanan and L. Battista for tissue collection.

## Funding

This work is supported by the EPFL Open Science Fund “Reproducible and Reusable Imaging Workflows” from the Imaging@EPFL initiative and the Swiss Data Science Center (SDSC). F.D.M. was supported by SNF (310030 179163/1). M.W. was supported by a generous donor represented by CARIGEST SA.

## Ethics Approval

Animal experiments were performed in accordance with protocol approved by the Service de la Consommation et des Affaires vétérinaires of Canton de Vaud (VD 1541.4 and VD 1865.3). NOD.Cg-Prkdcscid Il2rgtm1Wjl/SzJ mice (NSG) breeders were purchased from Jackson Laboratories.

## Consent to participate

The cantonal ethics committee approved the study on patient samples (183/10). Informed consent was obtained from all subjects.

## Availability of Data and Material

Archived images as at time of publication: DOI 10.5281/zenodo.3960270.

## Code Availability

The plugin source is accessible online and open-source:

https://github.com/Biomedical-Imaging-Group/SingleCellClassifier.

## Author Contributions

Conceptualization: Fabio De Martino.

Data Curation: Quentin Juppet, Fabio De Martino.

Formal Analysis: Quentin Juppet, Martin Weigert.

Funding Acquisition: Daniel Sage, Cathrin Brisken.

Investigation: Fabio De Martino.

Methodology Development: Quentin Juppet, Daniel Sage, Olivier Burri.

Project Administration: Fabio De Martino.

Resources: Fabio De Martino, Cathrin Brisken.

Software: Quentin Juppet, Daniel Sage.

Supervision: Fabio De Martino, Michaël Unser, Cathrin Brisken, Daniel Sage.

Writing - original draft: Fabio De Martino, Quentin Juppet, Martin Weigert, Olivier Burri.

Writing – Review & Editing: Fabio De Martino, Quentin Juppet, Martin Weigert, Michaël Unser, Cathrin Brisken, Daniel Sage.

## Conflicts of Interest/Competing Interests

The authors declare that they have no conflict of interest.

## Abbreviations

SCC: Single Cell Classifier
H&E: Haematoxylin and Eosin
HBECs: Human Breast Epithelial Cells
HR: Hormone Receptor
MIND: Mouse INtraDuctal
PDX: Patient-Derived Xenograft

## Supportive Information

**Sup. Fig 1.**
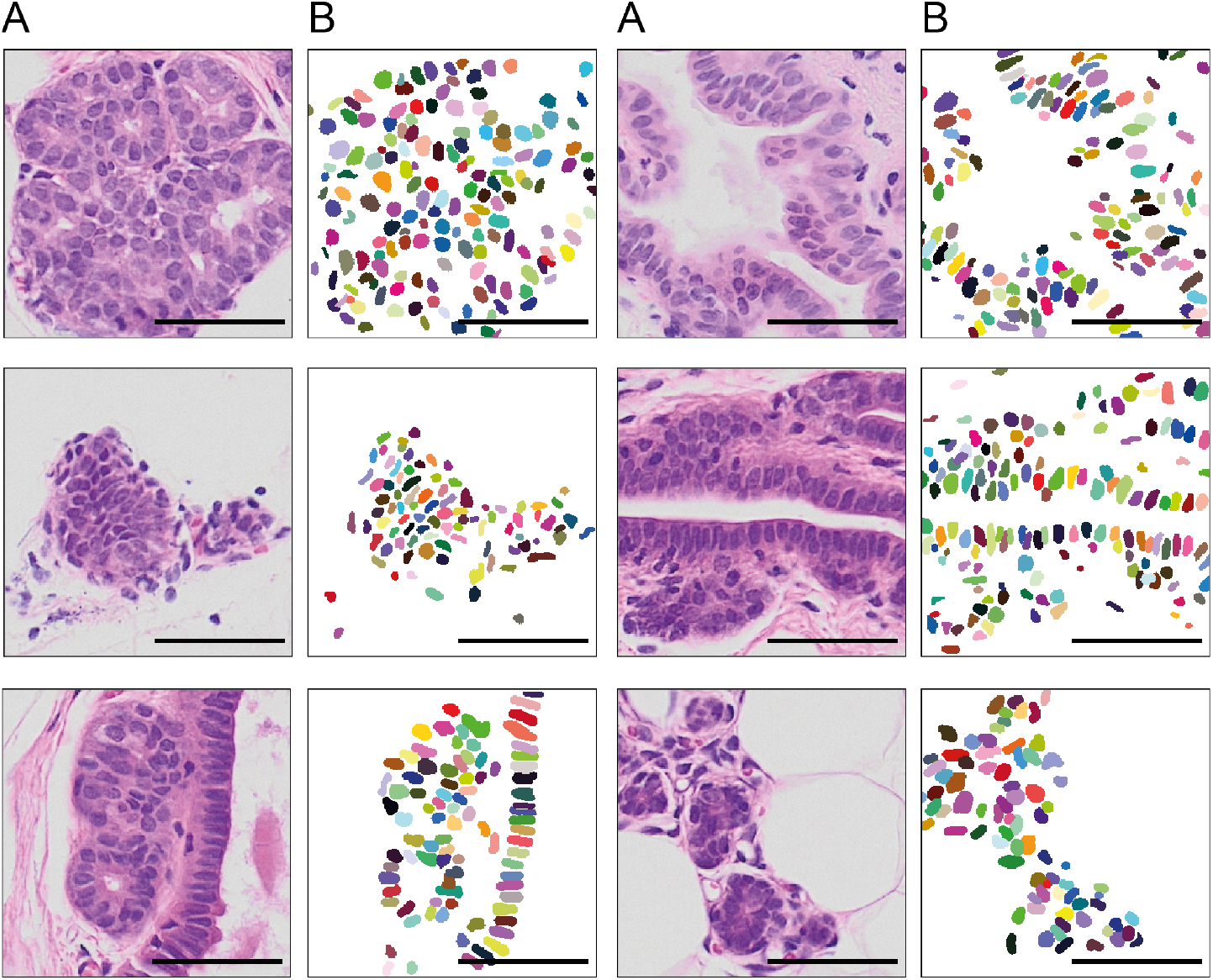
Example of the dataset used for the training of the StarDist model. A: The source images. B: The manual annotations of the nuclei associated with the source randomly colored for distinction purpose. Scale bars, 50 µm.

**Sup. Fig 2.**
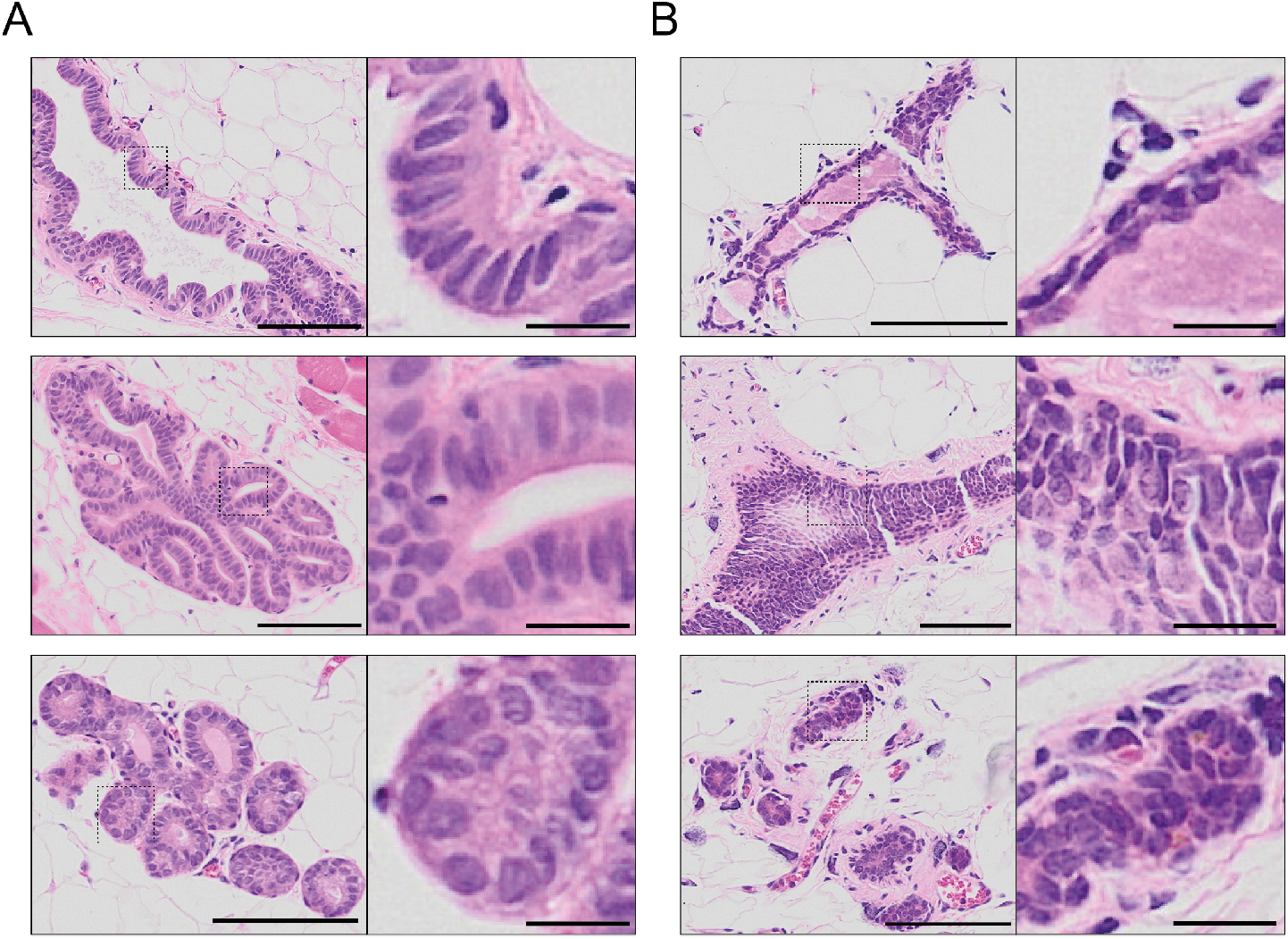
Example of the dataset used in the training of the classifier. A: Human labelled images. B: Mouse labelled images. Scale bars, 100 µm. Scale bars in magnified areas, 20 µm

## Notes

### Competing Interest Statement

The authors have declared no competing interest.

## References

1. Arrowsmith J. Phase II failures: 2008–2010. Nature Reviews. 2011; doi:10.1038/nrd3439.

2. Dimasi J, Reichert J, Feldman L, Malins A. Clinical Approval Success Rates for Investigational Cancer Drugs. Clinical pharmacology and therapeutics. 2013;94. doi:10.1038/clpt.2013.117.a

3. Dobrolecki L, Airhart S, Alferez D, Aparicio S, Behbod F, Bentires-Alj M, et al. Patient-derived xenograft (PDX) models in basic and translational breast cancer research. Cancer and Metastasis Reviews. 2016;35. doi:10.1007/s10555-016-9653-x.

4. Eirew P, Steif A, Khattra J, Ha G, Yap D, Farahani H, et al. Dynamics of genomic clones in breast cancer patient xenografts at single-cell resolution. Nature. 2014;518. doi:10.1038/nature13952.

5. Hidalgo M, Amant F, Biankin A, Budinská E, Byrne Phd A, Caldas C, et al. Patient-Derived Xenograft Models: An Emerging Platform for Translational Cancer Research. Cancer discovery. 2014;4:998–1013. doi:10.1158/2159-8290.CD-14-0001.

6. Haricharan S, Lei J, Ellis M. Mammary Ductal Environment Is Necessary for Faithful Maintenance of Estrogen Signaling in ER+ Breast Cancer. Cancer Cell. 2016;29:249–250. doi:10.1016/j.ccell.2016.02.017.

7. Kratz A, Ferrare M, Sluss P, Lewandrowski KB. Laboratory reference values. N Engl J Med. 2004;351:1548–1564.

8. Behbod F, Kittrell F, Machado H, Edwards D, Kerbawy S, Heestand J, et al. An intraductal human-in-mouse transplantation model mimics the subtypes of ductal carcinoma in situ. Breast cancer research: BCR. 2009;11:R66. doi:10.1186/bcr2358.

9. Sflomos G, Dormoy V, Metsalu T, Jeitziner R, Battista L, Scabia V, et al. A Preclinical Model for ERa-Positive Breast Cancer Points to the Epithelial Microenvironment as Determinant of Luminal Phenotype and Hormone Response. Cancer Cell. 2016;29:1–16. doi:10.1016/j.ccell.2016.02.002.

10. Siersbæk R, Scabia V, Nagarajan S, Chernukhin I, Papachristou EK, Broome R, et al. IL6/STAT3 Signaling Hijacks Estrogen Receptor α Enhancers to Drive Breast Cancer Metastasis. Cancer Cell. 2020;doi:https://doi.org/10.1016/j.ccell.2020.06.007.

11. Richard E, Grellety T, Velasco V, MacGrogan G, Bonnefoi H, Iggo R. The mammary ducts create a favourable microenvironment for xenografting of luminal and molecular apocrine breast tumours. The Journal of Pathology. 2016;240(3):256–261. doi:10.1002/path.4772.

12. Ataca D, Aouad P, Constantin C, Laszlo C, Beleut M, Shamseddin M, et al. The secreted protease Adamts18 links hormone action to activation of the mammary stem cell niche. Nature Communications. 2020;11(1):1571. doi:10.1038/s41467-020-15357-y.

13. Russell TD, Jindal S, Agunbiade S, Gao D, Troxell M, Borges VF, et al. Myoepithelial cell differentiation markers in ductal carcinoma in situ progression. The American journal of pathology. 2015;185(11):3076–3089. doi:10.1016/j.ajpath.2015.07.004.

14. Koch C, Kuske A, Joosse SA, Yigit G, Sflomos G, Thaler S, et al. Characterization of circulating breast cancer cells with tumorigenic and metastatic capacity. EMBO Molecular Medicine. 2020; p. e11908. doi:10.15252/emmm.201911908.

15. Danuser G. Computer Vision in Cell Biology. Cell. 2011;147:973–8. doi:10.1016/j.cell.2011.11.001.

16. McKinney S, Sieniek M, Godbole V, Godwin J, Antropova N, Ashrafian H, et al. International evaluation of an AI system for breast cancer screening. Nature. 2020;577:89–94. doi:10.1038/s41586-019-1799-6.

17. Janssens T, Antanas L, Derde S, Vanhorebeek I, Berghe G, Guiza F. CHARISMA: An integrated approach to automatic H&E-stained skeletal muscle cell segmentation using supervised learning and novel robust clump splitting. Medical image analysis. 2013;17:1206–1219. doi:10.1016/j.media.2013.07.007.

18. Chen H, Qi X, Yu L, Dou Q, Qin J, Heng PA. DCAN: Deep Contour-Aware Networks for Object Instance Segmentation from Histology Images. Medical Image Analysis. 2016;36. doi:10.1016/j.media.2016.11.004.

19. Salvi M, Molinari F. Multi-tissue and multi-scale approach for nuclei segmentation in H&E stained images. BioMedical Engineering OnLine. 2018;17. doi:10.1186/s12938-018-0518-0.

20. Schindelin J, Arganda-Carreras I, Frise E, Kaynig V, Longair M, Pietzsch T, et al. Fiji: an open-source platform for biological-image analysis. Nature Methods. 2012;doi:10.1038/nmeth.2019.

21. Bankhead P, Loughrey MB, Fernández JA, Dombrowski Y, McArt DG, Dunne PD, et al. QuPath: Open source software for digital pathology image analysis. Scientific Reports. 2017;doi:10.1038/s41598-017-17204-5.

22. Schmidt U, Weigert M, Broaddus C, Myers G. Cell Detection with Star-Convex Polygons. Medical Image Computing and Computer Assisted Intervention - MICCAI 2018 - 21st International Conference, Granada, Spain, September 16-20, 2018, Proceedings, Part II. 2018; p. 265–273. doi:10.1007/978-3-030-00934-2 30.

23. Weigert M, Schmidt U, Haase R, Sugawara K, Myers G. Star-convex Polyhedra for 3D Object Detection and Segmentation in Microscopy. The IEEE Winter Conference on Applications of Computer Vision (WACV). 2020;.

24. Haralick RM, Shanmugam K, Dinstein I. Textural Features for Image Classification. IEEE Transactions on Systems, Man, and Cybernetics. 1973;SMC-3(6):610–621.

25. Miyamoto E, Jr T. FAST CALCULATION OF HARALICK TEXTURE FEATURES. 2008;.

26. Chew LP. Constrained delaunay triangulations. Algorithmica. 1989; p. 97–108. doi:10.1007/BF01553881.

27. Lamouille S, Xu J, Derynck R. Molecular mechanisms of epithelial–mesenchymal transition. Nature Reviews Molecular Cell Biology. 2014;15:178–196. doi:10.1038/nrm3758.

28. Legland D, Arganda-Carreras I, Andrey P. MorphoLibJ: integrated library and plugins for mathematical morphology with ImageJ. Bioinformatics. 2016;32(22):3532–3534. doi:10.1093/bioinformatics/btw413.

29. Weigert M, Schmidt U, Boothe T, Móller A, Dibrov A, Jain A, et al. Content-aware image restoration: pushing the limits of fluorescence microscopy. Nature Methods. 2018;15(12):1090–1097. doi:10.1038/s41592-018-0216-7.

